# The most comprehensive annotation of the Krill transcriptome provides new insights for the study of physiological processes and environmental adaptation

**DOI:** 10.1101/2021.07.15.452476

**Authors:** Ilenia Urso, Alberto Biscontin, Davide Corso, Cristiano Bertolucci, Chiara Romualdi, Cristiano De Pittà, Bettina Meyer, Gabriele Sales

## Abstract

The krill species *Euphausia superba* plays a critical role in the food chain of the Antarctic ecosystem, as the abundance of its biomass affects trophic levels both below it and above. Major changes in climate conditions observed in the Antarctic Peninsula region in the last decades have already altered the distribution of the krill population and its reproductive dynamics. A deeper understanding of the adaptation capabilities of this species, and of the molecular mechanisms behind it are urgently needed. The availability of a large body of RNA-seq assays gave us the opportunity to extend the current knowledge of the krill transcriptome, considerably reducing errors and redundancies. Our study covered the entire developmental process, from larval stages to adult individuals, providing information of central relevance for ecological studies. Here we describe the KrillDB^2^ database, a resource combining the latest annotation of the krill transcriptome with a series of analyses specifically targeting genes and molecular processes relevant to krill physiology. KrillDB^2^ provides in a single resource the most complete collection of experimental data and bioinformatic annotations: it includes an extended catalog of krill genes; an atlas of their expression profiles over all RNA-seq datasets publicly available; a study of differential expression across multiple conditions such as developmental stages, geographical regions, seasons, and sexes. Finally, it provides initial indications about non-coding RNAs, a class of molecules whose contribute to krill physiology has never been reported before.

## Introduction

Antarctic krill *Euphausia superba* (hereafter krill) represents a widely distributed crustacean of the Southern Ocean and one of the world’s most abundant species with a total biomass between 100 and 500 million tonnes [1]. Due to its crucial ecological role in the Antarctic ecosystem, where it represents a link between apex predators and primary producers, several studies have been carried out over the years in order to characterize krill distribution [2, 3, 4], population dynamics and structuring [5, 6] and above all to understand its complex genetics [5, 7, 8, 9]. A sizable fraction of these studies focused on the DNA, specifically on mtDNA variation; however, the information available about krill genetics remains relatively modest. The difficulty in the progression of this kind of study mainly depends on the extraordinarily large krill genome size [10], which is more than 15 times larger than the human genome. This aspect largely complicates DNA sequencing, which is the reason why in recent years - together with the advances in high-throughput RNA-sequencing techniques - different krill transcriptome resources have been developed [11–16]. However, it was with the KrillDB project [17] that a detailed and advanced genetic resource was produced and made available to the community as an organized database. KrillDB is a web-based graphical interface with annotation results coming from the *de novo* reconstruction of krill transcriptome, assembling more than 360 million Illumina sequence reads.

Here we describe an update to KrillDB, now renamed KrillDB^2^ (available at the address https://krilldb2.bio.unipd.it/), specifically focusing on two aspects: the improvement of the quality and breadth of the krill transcriptome sequences previously reconstructed, thanks to the addition of an unprecedented amount of RNA-sequencing data; and, correspondingly, an increase in the amount of annotation information associated to each transcript. This improvement is made available through interactive graphs, images and downloadable files.

## Material and methods

### Krill collection

This study aims at covering the entire developmental process of krill. Therefore, we used samples coming from different developmental stages to cover the entire *E. superba* transcriptome, from larval to adult specimens. Specifically, adults included both male and female specimens, as well as summer and winter individuals and they also came from 3 different geographical regions: Lazarev Sea, South Georgia, and Bransfield Strait/South Orkney. The entire samples collection used to produce the new transcriptomic reference and carry out all downstream analysis is listed in **Table S1** (Supplementary Material).

### Transcriptome assembly strategy

#### Multiple independent *de novo* assemblies

The assembly of short (Illumina) reads to reconstruct the transcriptomes of non-model organisms has been subject to a considerable amount of research. Out of the many tools developed for this task, we selected the five which are arguably the most popular in the field: Trinity [18], BinPacker [19], rnaSPAdes [20], TransABySS [21] and IDBA-tran [22]. We summarized all the steps of the assembly reconstruction strategy, annotation process and downstream analyses in **Fig 1**.

**Fig 1.**
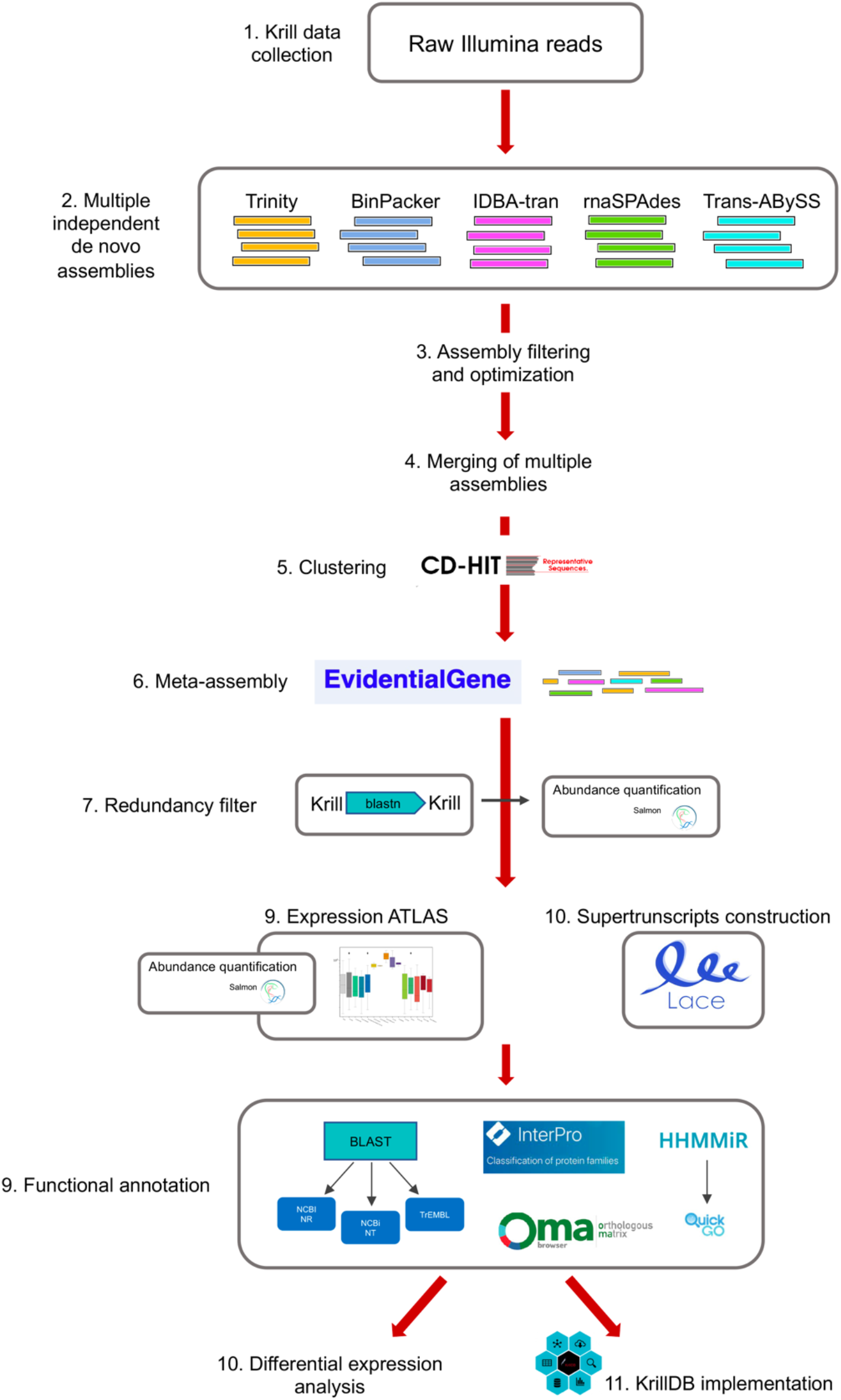
Workflow of the assembly process, annotation, database re-design and downstream analyses.

At first, we performed a separate transcriptome reconstruction with each of the tools listed above. We evaluated their respective advantages through a series of independent measures, such as: the total number of transcripts; %GC content; the average fragment length; the total number of bases; the N50 value; and finally, the results of the BUSCO analysis, which provides a measure of transcriptome completeness based on evolutionarily informed expectations of gene content from near-universal single-copy orthologs [23].

#### Assembly filtering and optimization

The raw sequencing data we used for the assemblies was obtained from different experiments and included both stranded (**Table S1** – Group 2) and unstranded libraries (**Table S1** – Group 1). As mixing these two types of libraries in a single assembly is not well supported, we decided to run each software twice: we thus generated a total of ten different *de novo* assemblies.

We used Trimmomatic [24] to remove adapter sequences and other artifacts from raw Illumina sequences. The, the quality of trimmed reads was checked with the program FastQC [25] (version 0.11.9). *De novo* transcriptome assembly was performed using specific parameters depending on the library type (the actual commands used are listed in **Table S2**, Supplementary Material).

Once assembled, a combination of two filtering steps was then applied to the newly reconstructed transcriptomes to discard artifacts and improve the assembly quality.

First, we estimated the abundances of all the transcripts reconstructed by each assembler using the Salmon software [26] (v. 1.4.0). Specifically, we used the following parameters were used: samples coming from unstranded library (**Table S1** – Group 1) were aligned using the options “-l ISR -1 --validateMappings”; samples coming from stranded library (**Table S1** – Group 2) were aligned using the options “-l IU --validateMappings”. Samples were grouped according to the main experimental conditions: (1) sex, with female and male levels; (2) geographical area, covering Bransfield Strait, South Georgia, South Orkney and Lazarev Sea; and (3) season, with summer and winter levels. Abundance estimates were imported in the R statistical environment using the *tximport* package [27] and we implemented a filter to keep only those transcripts showing an expression level of at least 1 transcript per million (TPM) within each of the three experimental conditions.

In a second step, we considered the results of all assemblers jointly, and we ran the “cd-hit-est” program [28] in order to cluster similar sequences and to produce a set of non-redundant representative transcripts. Specifically, we collapsed all sequences sharing 95% or more of their content, thus reducing the number of transcripts from 1,650,404 to 551,110.

#### Meta-assembly

The procedure described above was designed to identify near-duplicate sequences deriving from different software, but likely corresponding to the same biological transcript. As a further refinement, we were also interested in grouping resulting transcripts into units corresponding to genes. To this end, we relied on the EvidentialGene pipeline [29, 30]. We applied the “tr2aacds” tool which clusters transcripts and classifies them to identify the most likely coding sequence representing each gene. The software subdivides sequences into different categories, including primary transcript with alternates (main), primary without alternates (noclass), alternates with high and medium alignment to primary (althi1, althi, altmid) and partial (part) incomplete transcripts. A “coding potential” flag is also added, separating coding from non-coding sequences (see section “KrillDB^2^ Web Interface”). The meta-assembly thus obtained consisted in 274,840 putative transcripts, subdivided into 173,549 genes.

As these figures remained unrealistically high, we performed another round of analyses to identify redundant or mis-assembled sequences still appearing in our transcriptome. Here we used a combination of BLAST searches against known protein and nucleotide databases (NR, NT, TREMBL) and information deriving from full-length, experimentally validated transcripts from a previous study [31]. Results confirmed that the newly reconstructed transcriptome fully represented krill RNAs, but the large amount of input reads, together with the number of independent *de novo* assemblers, likely led to an inflation in the number of alternative splicing variants being reconstructed. Moreover, transcript alignments against BUSCO genes [23] and the *doubletime*, *cry1, shaggy* and *vrille* full-length transcripts from [31] highlighted the fact that multiple fragments of the same gene were incorrectly assembled as separate transfrags. To remove these artifacts, first we aligned all transcript sequences in our meta-assembly against each other using the *blastn* tool. We discarded all sequences already included in a longer transcript for more than the 90% of their length. This filter helped us remove 78,731 redundant sequences (29% of transcripts, overall). Then, we ran a new abundance quantification using Salmon and we discarded all transcripts with an average abundance below 0.1 TPM.

The combination of all the filters discussed above allowed us to reduce the number of transcripts to 151,585 and, correspondingly, that of genes to 85,905. Our approach discarded redundant genes, while retaining alternative transcripts with a sufficient level of uniqueness in their sequence. This was confirmed by the fact that although we removed almost 45% of the initially assembled transcripts, this filtering barely affected the average read mapping rate, which went from 89% (initial EvidentialGene output) to 88% (full filtering).

In order to enhance the interpretability of the transcriptome reconstruction, we also employed a SuperTranscripts analysis, on the basis of the workflow proposed by [32]. Specifically, we ran the Lace software (https://github.com/Oshlack/Lace) to reconstruct the block structure of each gene (see section “KrillDB^2^ Web Interface”).

### Functional Annotation

Assembled fragments were aligned against the NCBI NR (non-redundant) UniProtKB/TrEMBL protein databases and against the NCBI NT nucleotide collection (data downloaded on 22/04/2021). We also ran InterproScan (version 5.51-85.0) in order to search for known functional domains and to predict protein family membership. Results with an *e-value* greater than 1e-6 for proteins (*blastx*) or 1e-9 for nucleotides (*blastn*) were discarded.

Orthology inference was performed using the Orthologus MAtrix (OMA) standalone package [33] (https://omabrowser.org/standalone/) which relies on a complete catalog of orthologous genes among more than 2,300 genomes covering the entire tree of life. This analysis helped us identify, based on protein sequences, those krill transcripts showing an orthology relationship with genes from other species and which sets of genes derived from a single common ancestral gene at a given taxonomic range [34].

Finally, all krill transcripts were compared against the RNAcentral database (https://rnacentral.org/; https://doi.org/10.1093/nar/gkw1008) in order to identify any homology with the mature sequences of known microRNAs from other species.

### Expression Atlas

We used the final assembly described above to re-estimate transcript abundances over a wide range of RNAseq dataset (see **Table S1**) including:

- Larval krill at two different stages of development exposed to different CO_2_ conditions, coming from [17] (**Table S1** – Group 1)
- Adult krill (48 samples) coming from different geographical areas (Bransfield Strait, Lazarev Sea, South Georgia, South Orkney) and different seasons (summer and winter), divided into male and female specimens [35] (**Table S1** – Group 2)
- Adult krill exposed to three different temperatures – Low Temperature, Mid temperature, High Temperature (**Table S1** – Group 3)
- Adult krill divided into male and female specimens [36] (**Table S1** – Group 4)

Overall, these datasets include six experimental factors: geographical area, season, developmental stage, pCO_2_ exposure condition, sex and temperature. Newly computed transcript abundances and raw counts were imported using R (version 4.0.5) and the package *tximport* (version 1.18.0). Batch effect removal was performed using the *removeBatchEffect* function implemented in the *limma* package (version 3.46.0). The resulting count matrix of transcripts (rows) across samples (columns) was then converted to the transcripts per million (TPM) scale. Finally, results were summarized to the gene level using the *isoformToGeneExp* function (IsoformSwitchAnalyzeR version 1.12.0). The expression levels for each experimental condition are displayed in KrillDB^2^ as a barplot, as part of the webpage for each gene or transcript (see section “KrillDB^2^ Web Interface”).

### Differential Expression Analysis

Transcript-level abundances and estimated counts were summarized at the gene-level using the package *tximport*. Resulting counts were normalized to remove unwanted variation by means of the RUVg method [37]. Specifically, we performed a preliminary between-sample normalization (EDASeq, version 2.24.0) to adjust for sequencing depth. Following the workflow outlined in the RUVseq vignette, we identified a set of negative control genes with an FDR level larger than 0.8. We applied the RUVg method to estimate k=2 factors of unwanted variation and we included those in the design matrix for the final differential expression analysis, performed using the GLM method implemented by the edgeR software (version 3.32.1). All p-values were corrected using the Benjamini-Hochberg method.

### MicroRNAs

We also investigated the possibility that the new transcriptome included sequences corresponding to the precursors of krill microRNAs.

To this aim, we ran the HHMMiR software [38], which combines structural and sequence information to train a Hierarchical Hidden Markov Model for the identification of microRNA genes. We also performed a *blastn* search of all our assembled transcripts against the collection of miRBase (http://www.mirbase.org/) mature sequences. Results from these two analyses were combined: we collected all transcripts with a HHMMiR score below or equal to 0.71 and an alignment to a known mature microRNA with at most two mismatches. We then used the QuickGO tool (https://www.ebi.ac.uk/QuickGO/) to identify any potential association among our putatively identified microRNA precursors and GO categories.

### Opsin phylogeny

To identify novel opsin genes in krill, we manually examined manually the list of transcripts that were annotated as “opsin” by our automated pipeline. Furthermore, the entire krill transcriptome was aligned against a curated opsin dataset (including 996 visual and non-visual opsins [39]) using Blast+ (version 2.11.0). For genes with multiple alternative variants, we selected the longest transcript as a representative sequence. Secondary structure was assessed by the NCBI Conserved Domain Search (CDD database, May 2021). A phylogenetic tree was generated using the MUSCLE alignment tool and the Maximum Likelihood method (Dayhof substitution matrix and Nearest-Neighbor-Interchange method) as implemented in MEGA X (version 10.2.6, https://www.megasoftware.net/). New opsins were aligned against a curated invertebrate-only opsin data set [40], the previously cloned krill opsins [41], and the full-length onychopsin and arthropsin sequences available on the NCBI Protein database (May 2021, ncbi.nlm.nih.gov/protein). The tree was rooted using the human G protein-coupled receptor VIPR1 as an outgroup.

### Web Interface Implementation

The website was developed as a Python application based on the Flask framework. Data is stored in a PostgreSQL 12.8 database (http://www.postgresql.com). The sequences of the assembled transcripts and corresponding proteins are available for download as FASTA files. Gene and transcript pages have been updated with barplots implemented using the Seaborn Python library (version 0.11.1).

## Results

### Transcriptome Quality

We checked the quality of our transcriptome reconstruction quality step by step, starting from the 10 independent *de novo* assemblies, then evaluating the potential of merging all assemblies into a unique meta-assembly through EvidentialGene, and finally filtering the transcriptome for redundancy. All these results are summarized in **Fig 2**. As previously mentioned, the result of our reconstruction strategy was evaluated using different measures: the N50 statistics highlighted an increase in transfrags lengths at each step. Recent benchmarks, such as [42], have shown that, while reconstructing the transcriptome of a species, no single approach is uniformly superior at reconstructing the transcriptome of a species: the quality of each result is influenced by a number of factors, both technical (*k*-mer size, strategy for duplicate resolution) and biological (genome size, presence of contaminants). In our study we observed that, although a consistent number of sequences was removed through each step of the assembly, merging and filtering procedure, we didn’t encounter any decline in the quality described by the basic statistics of the reconstructed transcripts.

**Fig 2.**
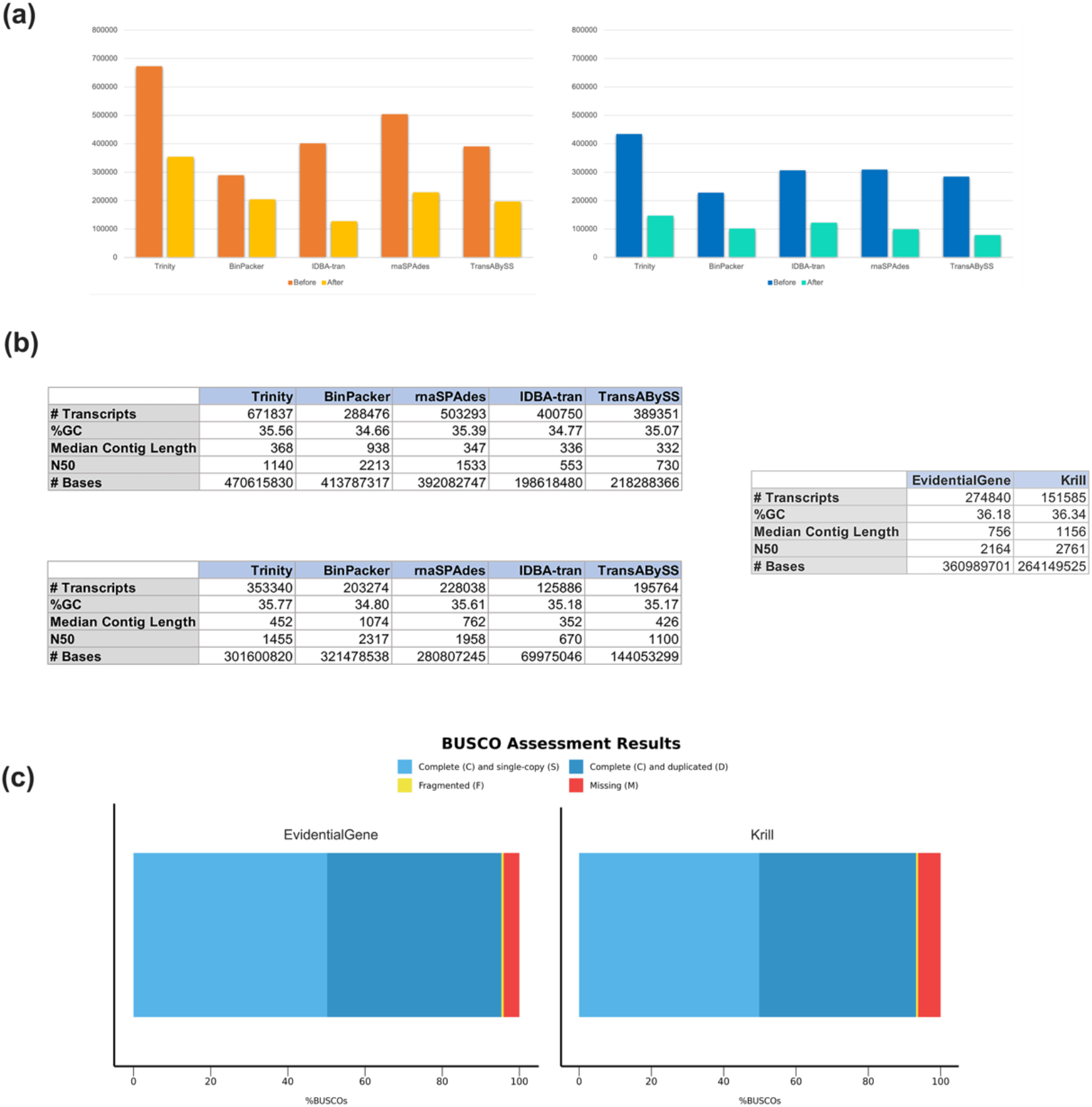
Transcriptome quality assessment results. Panel **(a)** shows the results of the first assembly filtering in terms of total number of transcripts. Quality measures computed at each assembly step are reported in panel **(b)**, from the five *de novo* assembly algorithms (top), after the first filtering process (bottom) and finally comparing the quality of the EvidentialGene meta-assembly and the final krill transcriptome after the redundancy filter (right). BUSCO assessment results **(c)** on EvidentialGene transcriptome (left) and krill transcriptome after last filter (right): the EvidentialGene transcriptome was characterized by 95.3% Complete sequences (50.2% single-copy, 45.1% duplicated), 0.6% Fragmented and 4.1% Missing sequences. The same analysis on the final krill transcriptome reconstruction produced 93.2% Complete transcripts (49.8% Single-copy, 43.4% Duplicated), 0.6% Fragmented and 6.2% Missing sequences.

We then explored the completeness of the krill transcriptome according to conserved ortholog content using BUSCO searching our sequences among all expected orthologs from Arthropoda phylum. This analysis confirms that our strategy for reducing redundancy did not affect transcriptome completeness: indeed, the fraction of complete single-copy essential genes drops by 2.1% while our strategy discards more than 44% redundant transfrags.

We finally compared our quality assessment results with those from previously released krill transcriptomes (**Table 1**). Our latest assembly significantly improves all the metrics we have discussed above and highlights the potential of the filtering strategy we have devised.

**Table 1.** Quality statistics of the previously released krill transcriptomes compared to the newly assembled KrillDB^2^. GenBank accession GFCS00000000.1 refers to the SuperbaSe krill transcriptome reference [43].

|  | GFCS000000000.1 | KrillDB | KrillDB<sup>2</sup> |
| --- | --- | --- | --- |
| **#Total Transcript** | 484080 | 133965 | 151585 |
| **Median Contig Length** | 439 | 683 | 1156 |
| **N50** | 1071 | 1294 | 2761 |
| **BUSCO - Complete** | 827 (81.6%) | 536 (52.9%) | 944 (93.2%) |

### Functional Classification

Results from the functional annotation analyses showed that 63,633 contigs matched at least one protein from the NCBI NR (non-redundant) collection, corresponding to about 42% of the total krill transcriptome, while 62,249 transfrags found a match among UniProtKB/TREMBL protein sequences (41% of the total). Furthermore, 22,071 krill transcripts (15% of the total) had significant matches with sequences in the NCBI NT nucleotide database. To classify transcripts by putative function, we performed a GO assignment. Specifically, 2,612 GO terms (corresponding to 13068 genes) were assigned: 1,128 of those (corresponding to 1178 genes) represented molecular functions; 1,099 terms (corresponding to 6991 genes) were linked to biological processes; 385 terms (corresponding to 4303 genes) represented cellular components.

### A case study on the discovery of opsin genes

To evaluate the gene discovery potential of the new assembly, we searched the transcriptome for novel members of the opsin family. Opsins are a group of light sensitive G protein-coupled receptors with seven transmembrane domains. 14 genes were annotated as putative opsins and the conserved domains analysis revealed that all of them possess the distinctive 7 α-helix transmembrane domain structure. The 8 previously cloned opsins [41] were all represented in KrillDB^2^ (sequence identity >90%; **Table S3**, Supplementary Material). The other 6 genes we identified can therefore be considered new putative opsins. Among those, we found 4 putative rhabdomeric opsins: *Es*Rh7 and *Es*Rh8, with 70% and 59 % of amino acid identity to *Es*Rh1a and *Es*Rh4, respectively; *Es*Rh9 and *Es*Rh10 showing high sequence identity (87% and 74%, respectively) to *Es*Rh5. Further, we identified 2 putative ancestral opsins: a non-visual arthropsin (*Es*Arthropsin), and an onychopsin (*Es*Onychopsin) with 70% and 49% of sequence identity with crustacean and onychophoran orthologous, respectively. Phylogenetic analysis (**Fig 3**) suggested that *Es*Rh7-10 are middle-wavelength-sensitive (MWS) rhabdomeric opsins, and further confirmed *Es*Arthropsin and *Es*Onychopsin annotation.

**Fig 3.**
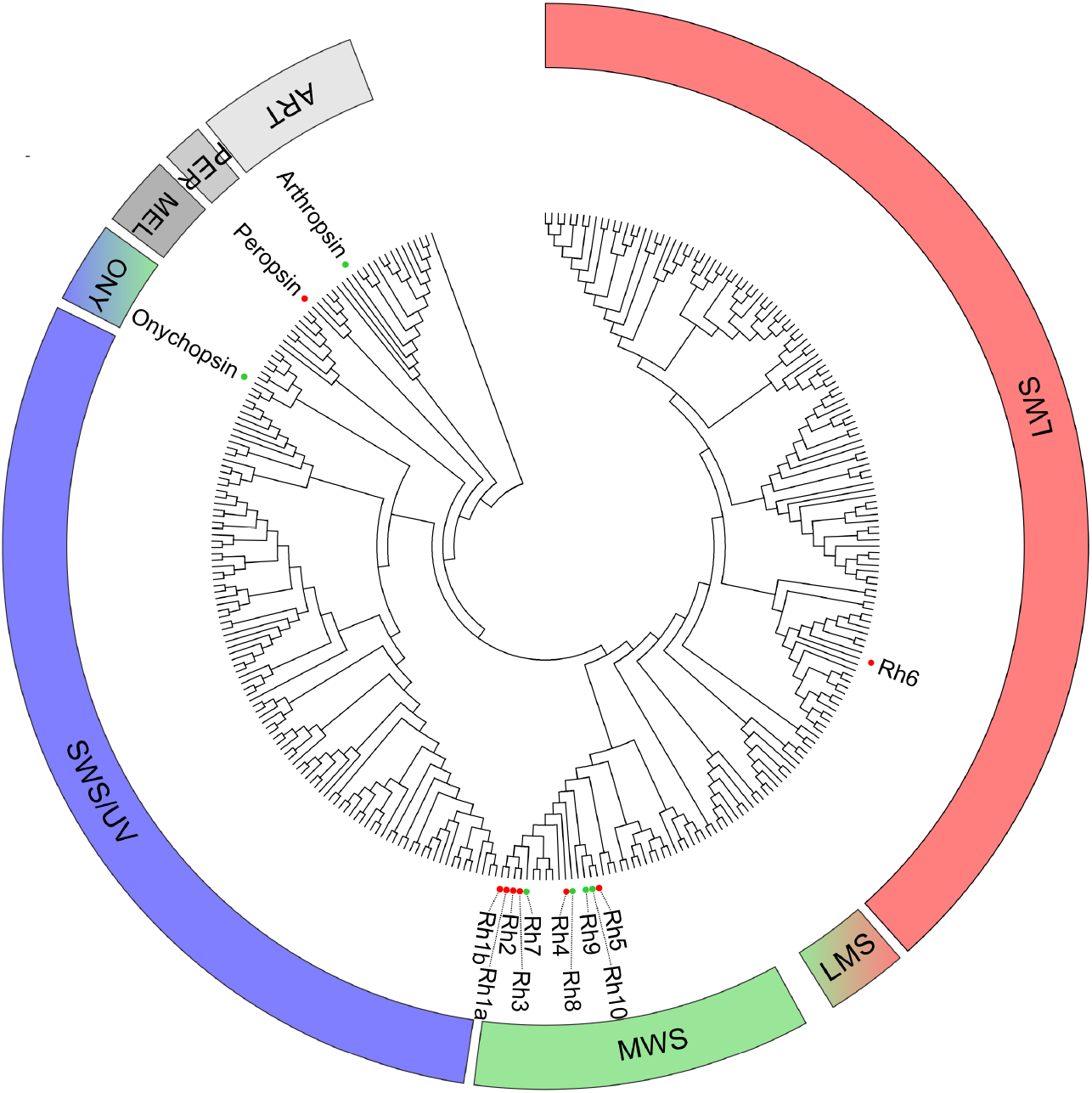
Phylogenetic relationships of *Euphausia superba* opsins shown as circular cladogram. Colored dots indicate krill opsins: red, previously cloned opsins; green, novel identified opsins. The spectral sensitivities of rhabdomeric opsin clades were inferred from the curated invertebrate-only opsin dataset proposed by DeLeo & Bracken-Grissom, 2020. Represented opsin classes: LWS, long-wavelenght-sensitive; LSM, long/middle-wavelenght-sensitive; MWS, middle-wavelenght-sensitive; SWS/UV, short/UV-wavelenght-sensitive; ONY, onychopsins; MEL, melanopsins; PER, peropsin; ART, arthropsin. Rectangular phylogram is reported in **Fig S1** (Supplementary Material).

### Differential Expression

The availability of a new assembly of the krill transcriptome, reconstructed collecting the largest amount of experimental data available thus far, suggested the possibility of performing a more detailed investigation of differential expression patterns. We decided to reanalyze the dataset from [35] to assess the possibility of identifying differentially expressed genes which were not detected in the original study due to the use of an older reference transcriptome [15].

Our design matrix for the model included all the independent factors (*season*, *area* and *sex*) and, in addition, the interaction between *area* and *season*, *sex* and *area*, *sex* and *season.*

In total 1,741 genes were found to be differentially expressed (DEG) among experimental conditions. They correspond to around 2% of the total reconstructed genes. In the previous work by [35] the same samples were quantified against a total of 58,581 contigs [15] producing a total of 1,654 DEGs. **Table 2** summarizes the list of contrasts that were performed, each one with the number of differentially expressed up and down regulated genes.

**Table 2.** List of contrast computed with total number of differentially expressed genes and numbers of up- and downregulated genes.

| Reference condition | Alternative condition | Sample group | # Total | # Upregulated | # Downregulated |
| --- | --- | --- | --- | --- | --- |
| Summer | Winter | Group 2 | 1195 | 1078 | 117 |
| Male | Femae | Group 2 | 14 | 7 | 7 |
| Male/Summer | Female/Winter | Group 2 | 12 | 6 | 6 |
| South Georgia | Lazarev Sea | Group 2 | 79 | 26 | 53 |
| South Georgia | Bransfield Strait-South Orkney | Group 2 | 28 | 6 | 22 |
| Lazarev Sea | Bransfield Strait-South Orkney | Group 2 | 17 | 13 | 4 |
| South Georgia/Male | Bransfield Strait-South Orkney/Female | Group 2 | 10 | 6 | 4 |
| South Georgia/Male | Lazarev Sea/Male | Group 2 | 19 | 8 | 11 |
| South Georgia/Summer | Bransfield Strait-South Orkney/Winter | Group 2 | 75 | 66 | 9 |
| Lazarev Sea/Summer | Bransfield Strait-South Orkney/Winter | Group 2 | 359 | 173 | 186 |
| South Georgia/Summer | Lazarev Sea/Summer | Group 2 | 188 | 150 | 38 |
| Lazarev Sea/Male | Bransfield Strait-South Orkney/Female | Group 2 | 20 | 10 | 10 |

1,195 DEGs were identified in the comparison between summer and winter specimens: 1,078 were up-regulated and 117 down-regulated. 396 of such DEGs had some form of functional annotation. In general, these results are in accordance with the discussion by Höring [35], which found that seasonal differences are predominant in comparison to regional ones. A summary of the DEGs is listed in **Table 3**. Complete tables of differentially expressed genes are downloadable on KrillDB^2^ (**Fig 4c**; https://krilldb2.bio.unipd.it/, Section “Differentially Expressed Genes (DEGs)”).

**Table 3.**
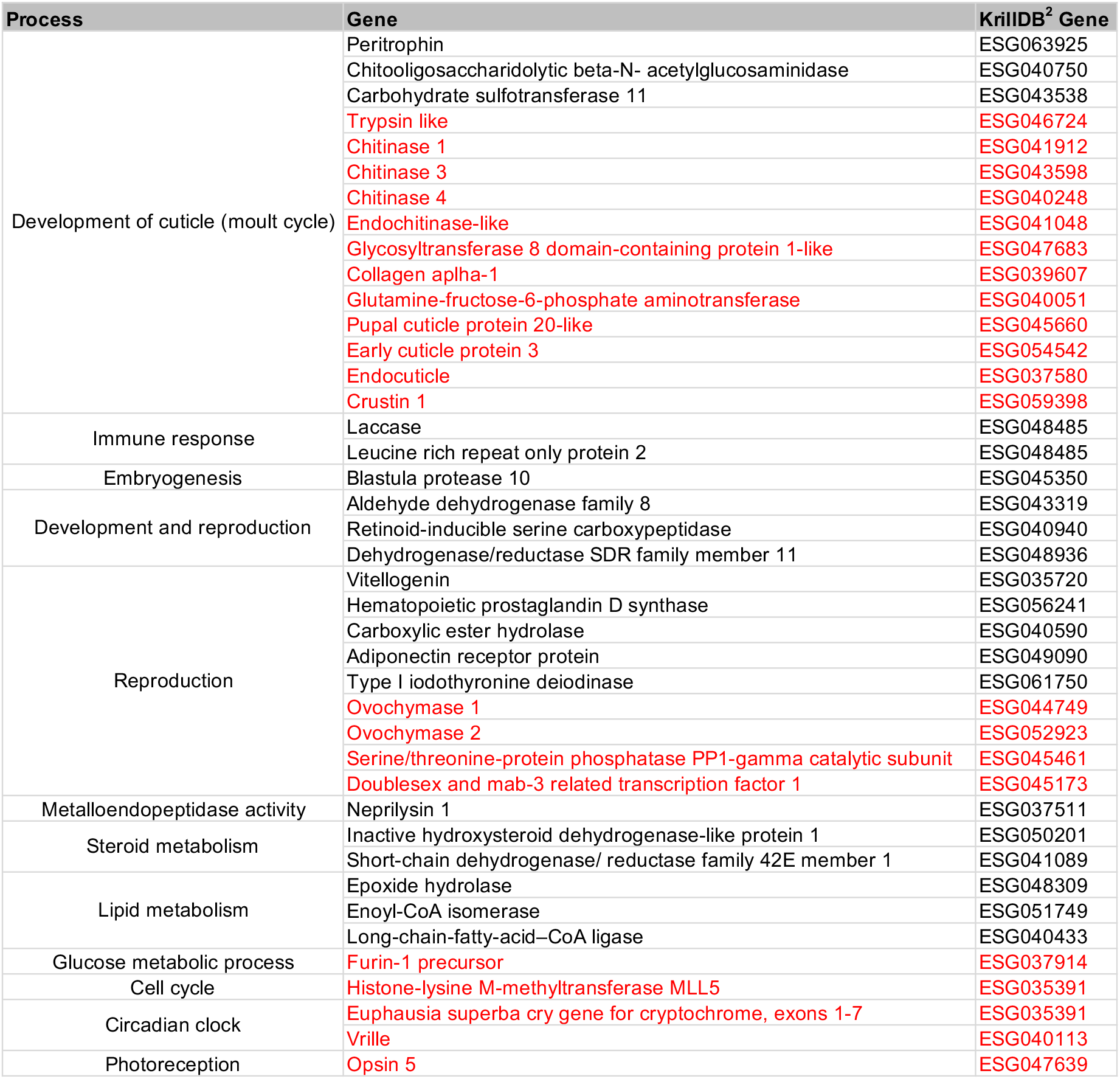
List of biologically relevant DEGs identified, starting from those already described by Höring et al [35]. Genes that were already found to be differentially expressed in the work by Höring are reported in black, while newly DEGs identified by our analysis are reported in red.

**Fig 4.**
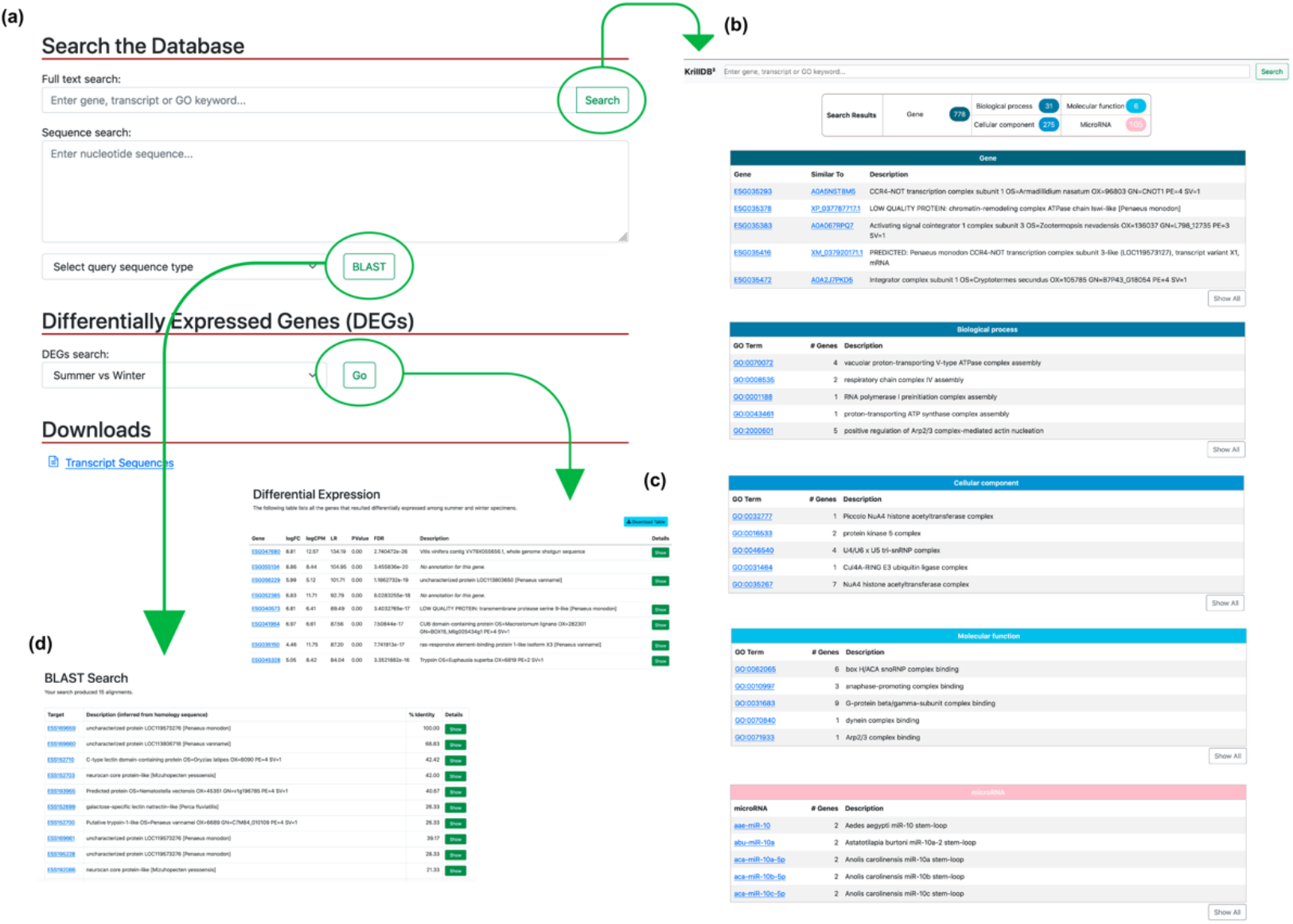
New search engine of KrillDB^2^. The homepage sections **(a)** with an example of the results of a fulltext-search **(b)**, of a blast-search **(d)** and the tables listing differentially expressed genes related to the contrast selected **(c)**.

### Summer vs Winter

We selected a series of genes among seasonal DEGs according to what has been already described in the literature. Höring et al. [35] previously identified and described 35 relevant DEGs involved in seasonal physiology and behavior: we recovered the same gene signature in our own analysis by comparing summer to winter samples. The majority of these DEGs appear to be involved in the development of cuticles (*chitine synthase*, *carbohydrate sulfotransferase 11*), lipid metabolism (*fatty acid synthase 2*, *enoyl-CoA ligase*), reproduction (*vitellogenin*, *hematopoietic prostaglandin D synthase*), metabolism of different hormones (*type 1 iodothyronine deiodinase*) and in the circadian clock (*cryptochrome*). Our results also include DEGs that were found to be involved in the moult cycle of krill in other studies [16]. Specifically, we identified a larger group of genes involved in the different stages of cuticle developmental process (*peritrophin-A domain*, *calcified cuticle protein*, *glycosyltransferase 8-domain containing protein 1*, *collagen alpha 1*, *glutamine-fructose 6 phosphate*), including proteins such as *cuticle protein-3,6,19.8, early cuticle protein, pupal cuticle protein, endocuticle structural glycoprotein, chitinase-3* and *chitinase-4*, the latter representing a group of chitinase which have been shown to be expressed predominantly in gut tissue during larval and/or adult stages in other arthropods and are proposed to be involved in digestion of chitin-containing substrates [44]. Finally, in addition to *trypsin* and *crustin 4* (immune-related gene, essential in early pre-moult stage when krill still have a soft cuticle to protect them from pathogen attack, as seen by Seear et al. [16]) we also identified *crustin-1,2,3,5* and *7*. All the reported genes were up-regulated in summer, the period in which growth take place and krill moult regularly.

Cuticle development genes were also identified as differentially expressed in the analysis of the interaction of multiple factors, in particular between male samples coming from South Georgia and female specimens coming from the area of Bransfield Strait-South Orkney (considered as a unique area since they are placed at similar latitudes). Strikingly, we also identified a pro-resilin gene, whose role in many insects consists in providing efficient energy storage, being upregulated in South Georgia male specimens.

### Interaction Effects

A number of relevant DEGs were found among specific interactions of regional and seasonal factors. In the comparison between krill samples in South Georgia in summer and individuals sampled in Bransfield Strait-South Orkney in winter we found genes, up-regulated in summer in South Georgia, that are related to reproductive activities, such as *doublesex* and *mab-3 related transcription factor*. The latter is a transcription factor crucial for sex determination and sexual differentiation which was already described in other arthropods [45]. Since no differentially expressed gene related to reproduction was found by Höring et al. [35] in the same comparisons, this suggests that the new krill transcriptome improves the interpretability of expression studies and the characterizations of krill samples.

Finally, the comparison between male individuals from the Lazarev Sea and female specimens from the Bransfield Strait-South Orkney showed additional DEGs involved in reproduction, such as *ovochymase 2*, usually highly expressed in female adults or eggs, *serine protease* and a *trypsin-like gene*. In particular, *trypsin-like genes* are usually thought to be digestive serine proteases, but previous works suggested that they can play other roles [46]; many trypsins show female or male-specific expression patterns and have been found exclusively expressed in males, as in our analysis, suggesting that they play a role in the reproductive processes.

The simultaneous presence of differentially expressed genes involved in different steps of the krill moulting cycle, in the reproductive process and in sexual maturation that appear to be differentially expressed in same comparisons is in accordance with what was already observed in krill [47] and other krill species [48]. In particular, there is evidence of a strong relation between the krill moulting process and its growth and sexual maturation during the year, which supports and confirms the reliability of our results in terms of genes involved in such krill life cycle steps.

### Identification of microRNA Precursors

In total we identified 261 krill transcripts with sequence homology to 644 known microRNAs from other species. 306 sequences were linked to at least one GO term, matching 54 krill transcripts (**Table S4**, Supplementary Material). Among them, we identified 5 putative microRNAs involved with changes in cellular metabolism (age-dependent general metabolic decline - GO:0001321, GO:0001323), as well as changes in the state or activity of cells (age-dependent response to oxidative stress - GO:0001306, GO:0001322, GO:0001324), 35 microRNAs involved in interleukin activity and production. We found 26 putative microRNAs likely involved in *ecdysteroidogenesis* (specifically GO:0042768), a process resulting in the production of ecdysteroids, moulting and sex hormones found in many arthropods. In addition, we found a microRNA involved in fused antrum stage (GO:0048165) which appears to be related in other species to oogenesis. We also identified 27 microRNAs related to *rhombomere* morphogenesis, formation and development (GO:0021661, GO:0021663, GO:0021570). These functions have been linked to the development of portions of the central nervous system in vertebrates, which share the same structure of those found in arthropod brains. Lastly, 26 krill sequences showed high similarity with 2 mature microRNA related to the formation of tectum (GO:0043676), which represents in arthropods and, specifically, crustaceans, the part of the brain acting as visual center.

### KrillDB^2^ Web Interface

The KrillDB website has been re-designed to include the new version of the transcriptome assembly. **Figs 4**, **5** and **6** collect images taken from the new main sections of the database. The integrated full-text search engine allows the user to search for a transcript ID, gene ID, GO term, a microRNA ID or any other free-form query. Results of full-text searches are now organized into several separate tables, each representing a different data source or biological aspect (**Fig 4b**). Results of GO term searches are summarized in a table reporting the related genes with corresponding domain (**Fig 5a**) or microRNA (**Fig 5b**) match and associated description. Both gene and transcript-centric pages have been extended with two new sections: “Orthology” and “Expression” (**Fig 6a**). The Orthology section summarizes the list of orthologous sequences coming from the OMA analysis, each one with the species it belongs to and the identity score.

**Fig 5.**
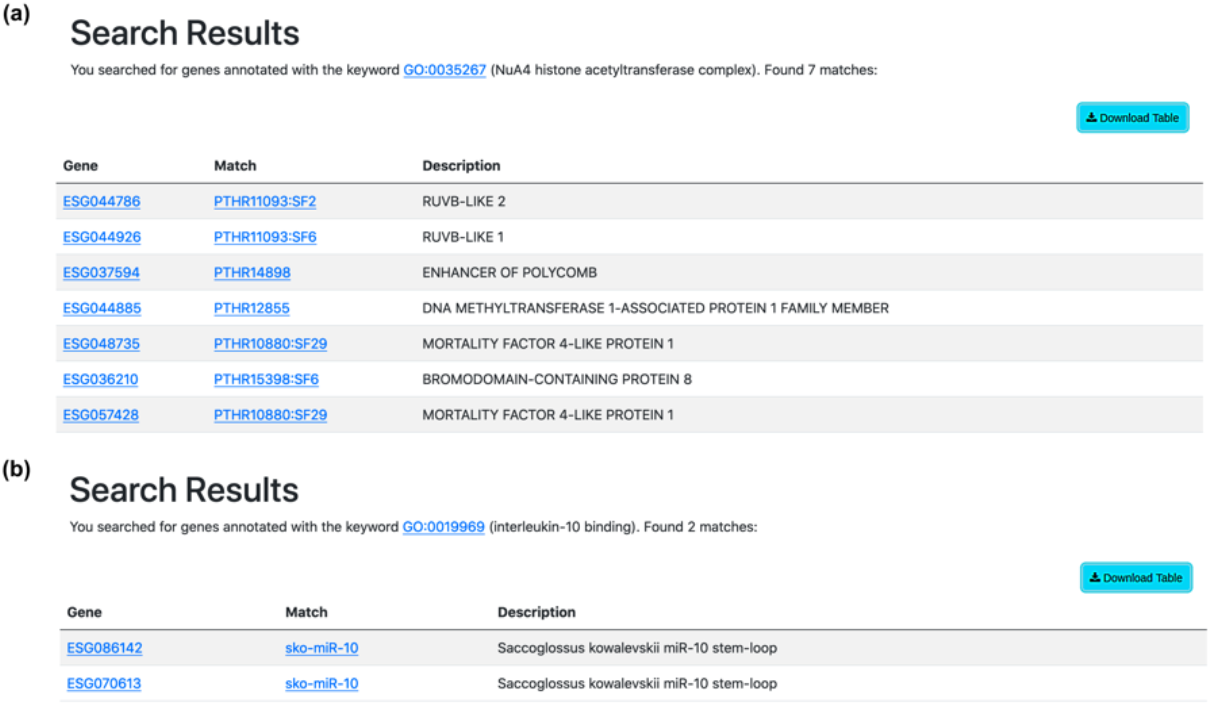
Table summarizing results of a GO term searc. An example of a GO term associated to genes matching different protein domains **(a)** and a GO term associated to genes matching a known mature microRNA **(b)**.

**Fig 6.**
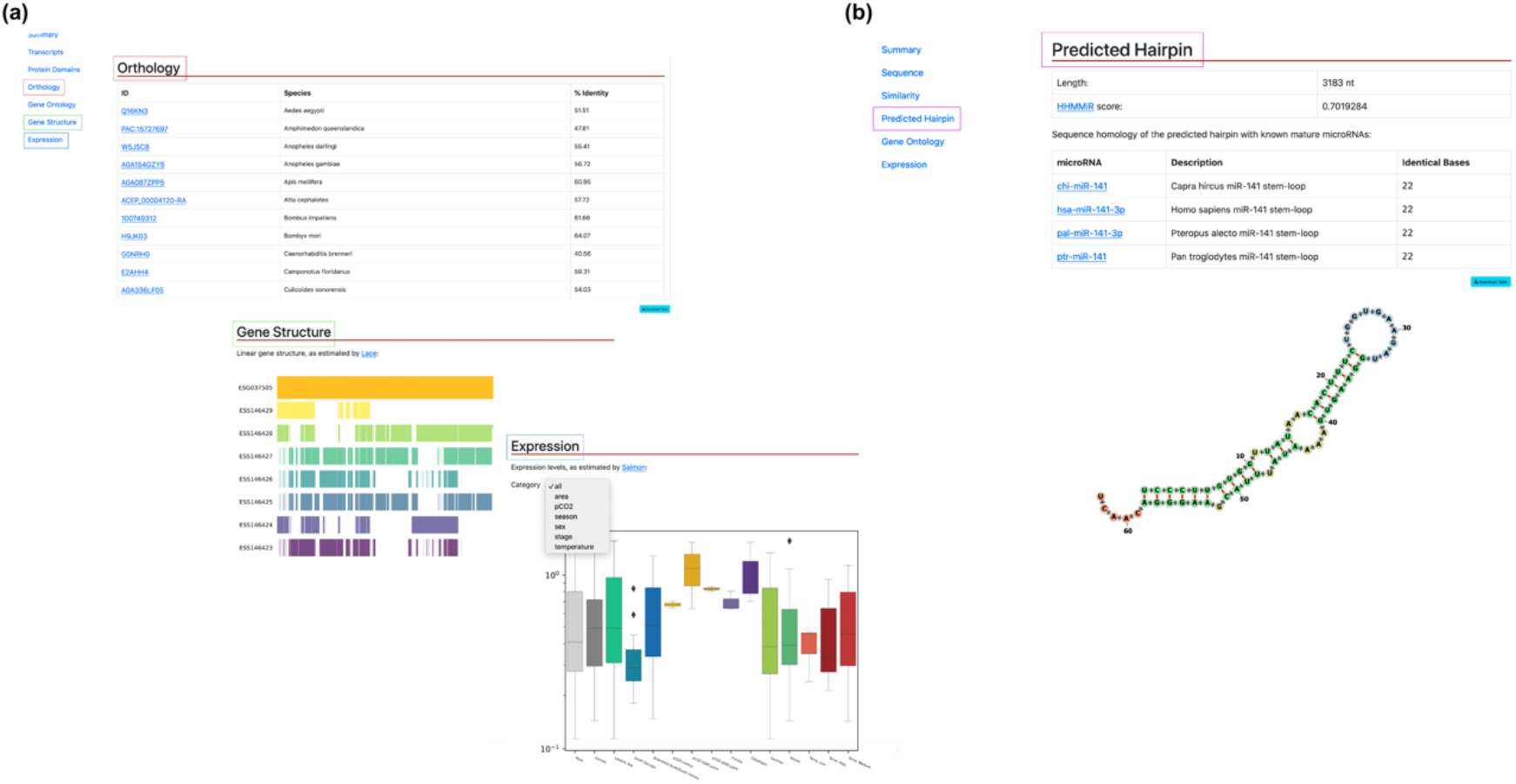
Additional sections in gene and transcript pages. The new sections in the gene-centric page show a table listing the orthologous sequences with their belonging species and the identity score, a visualization of the gene structure as estimated by Lace software and a boxplot coming from Expression Atlas analyses **(a)**. Both Orthology and Expression section are integrated also in the transcript-centric page. When a transcript is annotated as a putative microRNA, a “Predicted Hairpin” section displays a visualization of the hairpin predicted secondary structure and tables showing the alignment length, the HHMMiR score and the list of mature microRNAs matching **(b)**.

The “Expression” section shows a barplot representing abundances estimates obtained from Salmon. An additional section, called “Gene Structure” (**Fig 6a**), was added to the gene page on the basis of the results coming from the SuperTranscript analysis. Specifically, we modified the STViewer.py Python script (from Lace), optimizing and adapting it to our own data and database structure, in order to produce a visualization of each gene with its transcripts. Since Lace relies on the construction of a single directed splice graph and it is not able to compute it for complex clusters with more than 30 splicing variants, this section is available for a selection of genes only.

The new KrillDB^2^ release includes completely updated transcript and gene identifiers. However, the user searching for a retired ID is automatically redirected to the page describing the newest definition of the appropriate transcript or gene.

The KrillDB^2^ homepage now includes two additional sections (**Fig 4a**): one is represented by the possibility to perform a BLAST search. Any nucleotide or protein sequence (*query*) can be aligned against krill sequences stored in the database. Results are summarized in a table containing information about the krill transcripts (*target*) that matched with the user’s query, and the e-value corresponding to the alignment. The other new section, called “Differentially Expressed Genes”, allows the user to browse all the tables listing the genes that were found to be differentially expressed among the conditions we have described above. A drop-down menu gives access to the different comparisons; DEG tables (**Fig 4c**) list for each gene its log fold-change, p- and FDR values as estimated by edgeR. Moreover, each gene is linked to a functional description (if available) inferred from sequence homology searches.

Information about krill transcripts that showed homology with an annotated microRNA is available in the section “Predicted Hairpin” (**Fig 6b**). It contains a summary table with details about the hairpin length and the similarity score (as estimated by HHMMiR), followed by full listing of all the corresponding mature microRNAs (including links to their miRBase page). In addition, an image displaying the predicted secondary structure of the hairpin is included (computed by the “fornac” visualization software from the ViennaRNA suite).

## Discussion

The availability of a large amount of public RNA-seq data capturing krill transcripts has given us the possibility to re-assemble its transcriptome and to significantly extend its annotation. We have now covered the entire developmental process of this species and included in our analysis individuals belonging to different seasons and affected by different environmental conditions. KrillDB^2^ provides the most complete source of information about the krill transcriptome and will offer a reliable starting point development of novel ecological studies. As shown in **Table 1**, the analysis of the quality of previously released krill transcriptome in comparison to the newly assembled KrillDB^2^ confirmed how the strategy applied did not produce any loss in terms of quality, although a consistent number of transcripts was removed. The quality metrics, in contrast, were improved both in terms of N50 statistics and transcriptome completeness: the fraction of complete single-copy essential genes reached the 93.2%.

The differential expression analysis we have performed highlights the importance of specific processes in the complex krill life cycle and in its adaptation capability to the harsh Antarctic environment.

The identification of 6 novel putative opsin sequences almost double the eight that were previously cloned, demonstrating a significant improvement in the gene discovery potential of this new version of krill transcriptome. The finding of four novel MWS rhabdomeric opsins, an onychopsin, and a non-visual arthropsin further enrich the opsin repertoire of *E. superba* shedding light on a complex photoreception system able to coordinate the physiological and behavioral responses to the extreme daily (diel vertical migration) and seasonal changes in photoperiod and spectral composition. Arthropsins are rhabdomeric non-visual opsins and its clade is the sister group of the bilaterian rhabdomeric opsins [49, 50]. It was first discovered in the crustacean *Daphnia pulex* and subsequently in other arthropods, onychophoran, molluscs, anallids and flatworms [49–53]. Of relevance is the identification of an onychopsin which has been suggested to be the common ancestor of *Panarthropoda* visual opsins [49], and possibly sensitive to wavelength from UV to green light [54]. *Es*Onychopsin could represent the short-wavelength sensitive opsin (SWS/UV) which we have long been searching for. Indeed, the absence of a SWS/UV opsin was truly unexpected in an organism that shows daily vertical migration reaching depth beyond the 30 m, where only short wavelength light can penetrate.

Finally, KrillDB^2^ includes the first evidence of the role of non-coding RNAs in krill. Although this is just a preliminary analysis, the results we have described already hint at a role of microRNAs in defining the adaptive capabilities of this species to the Antarctic environment. This represents a promising starting point for the study of non-coding RNAs in the Antarctic krill and in other species belonging to the same family.

## Supporting information

Suppermentary Material

## Acknowledgments

The position of Ilenia Urso was supported by the Helmholtz Virtual Institute “PolarTime”: Biological timing in a changing marine environment - clocks and rhythms in polar pelagic organisms (VH-VI-500), headed by Bettina Meyer. Alberto Biscontin was funded by the “Programma Nazionale di Ricerche in Antartide – PNRA” (grant 2016_00225) and by the Promega Corporation 2019 Real-Time PCR Grant Program. We would also like to acknowledge the CAPRI initiative (Calcolo ad Alte Prestazioni per la Ricerca e l’Innovazione”, University of Padova Strategic Research Infrastructure Grant 2017) for the technical support and the HPC resources we have used for the analyses. Cristiano Bertolucci was supported by the “Programma Nazionale di Ricerche in Antartide – PNRA” (grant 2016_00225) and by the University of Ferrara research grant (FIR2020 and FAR2021).

## Data Accessibility

Data used for the krill transcriptome reconstruction and for the generation of the Expression Atlas was downloaded from the NCBI Short Read Archive, under accessions: PRJEB30084, PRJNA362526, PRJEB30084, PRJNA362526 and PRJNA640244.

## Author Contributions

IU, CDP, BM and GS conceived the study. IU and DC performed the analyses. IU, AB, BM and GS wrote the manuscript. CB, CR and CDP advised on data analysis and reviewed the text.

## References

1. Nicol, S., & Endo, Y. (1997). Krill fisheries of the world (No. 367). Food & Agriculture Org..

2. Atkinson, A., Siegel, V., Pakhomov, E. A., Rothery, P., Loeb, V., Ross, R. M., … & Fleming, A. H. Atkinson, A., Siegel, V., Pakhomov, E. A., Rothery, P., Loeb, V., Ross, R. M., … & Fleming, A. H. (2008). Oceanic circumpolar habitats of Antarctic krill. Marine Ecology Progress Series, 362, 1–23., 2008. doi: org/10.3354/meps07498

3. Hofmann, E. E., & Murphy, E. J. (2004). Advection, krill, and Antarctic marine ecosystems. Antarctic Science, 16(4). doi: 10.1017/s0954102004002275

4. Siegel, V. (2005). Distribution and population dynamics of Euphausia superba: summary of recent findings. Polar Biology, 29(1), 1–22. doi: 10.1007/s00300-005-0058-5

5. Bortolotto, E., Bucklin, A., Mezzavilla, M., Zane, L., & Patarnello, T. (2011). Gone with the currents: lack of genetic differentiation at the circum-continental scale in the Antarctic krill Euphausia superba. BMC genetics, 12(1), 1–18. doi: 10.1186/1471-2156-12-32

6. Valentine, J. W., & Ayala, F. J. (1976). Genetic variability in krill. Proceedings of the National Academy of Sciences, 73(2), 658–660. doi: 10.1073/pnas.73.2.658.

7. Batta-Lona, P. G., Bucklin, A., Wiebe, P. H., Patarnello, T., & Copley, N. J. (2011). Population genetic variation of the Southern Ocean krill, Euphausia superba, in the Western Antarctic Peninsula region based on mitochondrial single nucleotide polymorphisms (SNPs). Deep Sea Research Part II: Topical Studies in Oceanography, 58(13-16), 1652–1661. doi: 10.1016/j.dsr2.2010.11.017

8. Goodall-Copestake, W. P., Perez-Espona, S., Clark, M. S., Murphy, E. J., Seear, P. J., & Tarling, G. A. (2010). Swarms of diversity at the gene cox1 in Antarctic krill. Heredity, 104(5), 513–518. doi: 10.1038/hdy.2009.188

9. Zane, L., Ostellari, L., Maccatrozzo, L., Bargelloni, L., Battaglia, B., & Patarnello, T. (1998). Molecular evidence for genetic subdivision of Antarctic krill (Euphausia superba Dana) populations. Proceedings of the Royal Society of London. Series B: Biological Sciences, 265(1413), 2387–2391. doi: 10.1098/rspb.1998.0588

10. Jeffery, N. W. (2012). The first genome size estimates for six species of krill (Malacostraca, Euphausiidae): large genomes at the north and south poles. Polar Biology, 35(6), 959–962. doi: 10.1007/s00300-011-1137-4

11. Clark, M. S., Thorne, M. A., Toullec, J. Y., Meng, Y., Peck, L. S., & Moore, S. (2011). Antarctic krill 454 pyrosequencing reveals chaperone and stress transcriptome. PLos one, 6(1), e15919. doi: 10.1371/journal.pone.0015919

12. De Pittà, C., Bertolucci, C., Mazzotta, G. M., Bernante, F., Rizzo, G., De Nardi, B., … & Costa, R. (2008). Systematic sequencing of mRNA from the Antarctic krill (Euphausia superba) and first tissue specific transcriptional signature. BMC genomics, 9(1), 1–14. doi: 10.1186/1471-2164-9-45

13. De Pittà, C., Biscontin, A., Albiero, A., Sales, G., Millino, C., Mazzotta, G. M., … & Costa, R. (2013). The Antarctic krill Euphausia superba shows diurnal cycles of transcription under natural conditions. PLoS One, 8(7), e68652. doi: 10.1371/journal.pone.0068652

14. Martins, M. J. F., Lago-Leston, A., Anjos, A., Duarte, C. M., Agusti, S., Serrão, E. A., & Pearson, G. A. (2015). A transcriptome resource for Antarctic krill (Euphausia superba Dana) exposed to short-term stress. Marine genomics, 23, 45–47. doi: 10.1016/j.margen.2015.04.008

15. Meyer, B., Martini, P., Biscontin, A., De Pittà, C., Romualdi, C., Teschke, M., … & Kawaguchi, S. (2015). Pyrosequencing and de novo assembly of Antarctic krill (Euphausia superba) transcriptome to study the adaptability of krill to climate-induced environmental changes. Molecular ecology resources, 15(6), 1460–1471. doi: 10.1111/1755-0998.12408

16. Seear, P. J., Tarling, G. A., Burns, G., Goodall-Copestake, W. P., Gaten, E., Özkaya, Ö., & Rosato, E. (2010). Differential gene expression during the moult cycle of Antarctic krill (Euphausia superba). BMC genomics, 11(1), 1–13. doi: 10.1186/1471-2164-11-582

17. Sales, G., Deagle, B. E., Calura, E., Martini, P., Biscontin, A., De Pittà, C., … & Jarman, S. (2017). KrillDB: A de novo transcriptome database for the Antarctic krill (Euphausia superba). PLoS One, 12(2), e0171908. doi: 10.1371/journal.pone.0171908

18. Grabherr, M. G., Haas, B. J., Yassour, M., Levin, J. Z., Thompson, D. A., Amit, I., … & Regev, A. (2011). Trinity: reconstructing a full-length transcriptome without a genome from RNA-Seq data. Nature biotechnology, 29(7), 644. doi: 10.1038%2Fnbt.1883

19. Liu, J., Li, G., Chang, Z., Yu, T., Liu, B., McMullen, R., … & Huang, X. (2016). BinPacker: packing-based de novo transcriptome assembly from RNA-seq data. PLoS computational biology, 12(2), e1004772. doi: 10.1371/journal.pcbi.1004772

20. Bushmanova, E., Antipov, D., Lapidus, A., & Prjibelski, A. D. (2019). rnaSPAdes: a de novo transcriptome assembler and its application to RNA-Seq data. GigaScience, 8(9), giz100. doi: 10.1093/gigascience/giz100

21. Zhao, Q. Y., Wang, Y., Kong, Y. M., Luo, D., Li, X., & Hao, P. (2011, December). Optimizing de novo transcriptome assembly from short-read RNA-Seq data: a comparative study. In BMC bioinformatics (Vol. 12, No. 14, pp. 1–12). BioMed Central. doi: 10.1186/1471-2105-12-S14-S2

22. Peng, Y., Leung, H. C., Yiu, S. M., Lv, M. J., Zhu, X. G., & Chin, F. Y. (2013). IDBA-tran: a more robust de novo de Bruijn graph assembler for transcriptomes with uneven expression levels. Bioinformatics, 29(13), i326–i334. doi: 10.1093/bioinformatics/btt219

23. Simão, F. A., Waterhouse, R. M., Ioannidis, P., Kriventseva, E. V., & Zdobnov, E. M. (2015). BUSCO: assessing genome assembly and annotation completeness with single-copy orthologs. Bioinformatics, 31(19), 3210–3212. doi: 10.1093/bioinformatics/btv351

24. Bolger, A. M., Lohse, M., & Usadel, B. (2014). Trimmomatic: a flexible trimmer for Illumina sequence data. Bioinformatics, 30(15), 2114–2120. doi: 10.1093/bioinformatics/btu170

25. Andrews, S. (2017). FastQC: a quality control tool for high throughput sequence data. 2010.

26. Patro, R., Duggal, G., Love, M. I., Irizarry, R. A., & Kingsford, C. (2017). Salmon provides fast and bias-aware quantification of transcript expression. Nature methods, 14(4), 417–419. doi: 10.1038/nmeth.4197

27. Love, M. I., Soneson, C., & Robinson, M. D. (2017). Importing transcript abundance datasets with tximport. Dim Txi. Inf. Rep. Sample1, 1, 5.

28. Li, W., & Godzik, A. (2006). Cd-hit: a fast program for clustering and comparing large sets of protein or nucleotide sequences. Bioinformatics, 22(13), 1658–1659. doi: 10.1093/bioinformatics/btl158

29. Gilbert, D. G. (2019). Genes of the pig, Sus scrofa, reconstructed with EvidentialGene. PeerJ, 7, e6374. doi: 10.7717/peerj.6374

30. Gilbert, D. G. (2019). Longest protein, longest transcript or most expression, for accurate gene reconstruction of transcriptomes? bioRxiv, 829184. doi: 10.1101/829184

31. Biscontin, A., Wallach, T., Sales, G., Grudziecki, A., Janke, L., Sartori, E., … & Costa, R. (2017). Functional characterization of the circadian clock in the Antarctic krill, Euphausia superba. Scientific reports, 7(1), 1–13. doi: 10.1038/s41598-017-18009-2

32. Davidson, N. M., Hawkins, A. D., & Oshlack, A. (2017). SuperTranscripts: a data driven reference for analysis and visualization of transcriptomes. Genome biology, 18(1), 1–10. doi: 10.5281/zenodo.830594

33. Altenhoff, A. M., Levy, J., Zarowiecki, M., Tomiczek, B., Vesztrocy, A. W., Dalquen, D. A., … & Dessimoz, C. (2019). OMA standalone: orthology inference among public and custom genomes and transcriptomes. Genome research, 29(7), 1152–1163. doi: 10.1101/gr.243212.118

34. Altenhoff, A. M., Gil, M., Gonnet, G. H., & Dessimoz, C. (2013). Inferring hierarchical orthologous groups from orthologous gene pairs. PloS one, 8(1), e53786. doi: 10.1371/journal.pone.0053786

35. Höring, F., Biscontin, A., Harms, L., Sales, G., Reiss, C. S., De Pittà, C., & Meyer, B. (2021). Seasonal gene expression profiling of Antarctic krill in three different latitudinal regions. Marine Genomics, 56, 100806. doi: 10.1016/j.margen.2020.100806

36. Suter, L., Polanowski, A. M., King, R., Romualdi, C., Sales, G., Kawaguchi, S., … & Deagle, B. E. (2019). Sex identification from distinctive gene expression patterns in Antarctic krill (Euphausia superba). Polar Biology, 42(12), 2205–2217. doi: 10.1007/s00300-019-02592-3

37. Risso, D., & Course, I. B. S. (2015). RNA-seq Normalization and Batch Effect Removal.

38. Kadri, S., Hinman, V., & Benos, P. V. (2009). HHMMiR: efficient de novo prediction of microRNAs using hierarchical hidden Markov models. BMC bioinformatics, 10(1), 1–12. doi: 10.1186/1471-2105-10-S1-S35

39. Henze, M. J., & Oakley, T. H. (2015). The dynamic evolutionary history of pancrustacean eyes and opsins. Integrative and comparative biology, 55(5), 830–842. doi: 10.1093/icb/icv100

40. DeLeo, D. M., & Bracken-Grissom, H. D. (2020). Illuminating the impact of diel vertical migration on visual gene expression in deep-sea shrimp. Molecular Ecology, 29(18), 3494–3510. doi: 10.1111/mec.15570

41. Biscontin, A., Frigato, E., Sales, G., Mazzotta, G. M., Teschke, M., De Pittà, C., … & Bertolucci, C. (2016). The opsin repertoire of the Antarctic krill Euphausia superba. Marine genomics, 29, 61–68. doi: 10.1016/j.margen.2016.04.010

42. Hölzer, M., & Marz, M. (2019). De novo transcriptome assembly: A comprehensive cross-species comparison of short-read RNA-Seq assemblers. GigaScience, 8(5), giz039. doi: 10.1093/gigascience/giz039

43. Hunt, B. J., Özkaya, Ö., Davies, N. J., Gaten, E., Seear, P., Kyriacou, C. P., Tarling G., Rosato, E. (2017). The Euphausia superba transcriptome database, Superba SE: An online, open resource for researchers. Ecology and evolution, 7(16), 6060–6077. doi: 10.1002/ece3.3168

44. Khajuria, C., Buschman, L. L., Chen, M. S., Muthukrishnan, S., & Zhu, K. Y. (2010). A gut-specific chitinase gene essential for regulation of chitin content of peritrophic matrix and growth of Ostrinia nubilalis larvae. Insect biochemistry and molecular biology, 40(8), 621–629. doi: 10.1016/j.ibmb.2010.06.003

45. Jia, L. Y., Chen, L., Keller, L., Wang, J., Xiao, J. H., & Huang, D. W. (2018). Doublesex evolution is correlated with social complexity in ants. Genome biology and evolution, 10(12), 3230–3242. doi: 10.1093/gbe/evy250

46. Bao, Y. Y., Qin, X., Yu, B., Chen, L. B., Wang, Z. C., & Zhang, C. X. (2014). Genomic insights into the serine protease gene family and expression profile analysis in the planthopper, Nilaparvata lugens. BMC genomics, 15(1), 1–17. doi: 10.1186/1471-2164-15-507

47. Buchholz, F., Watkins, J. L., Priddle, J., Morris, D. J., & Ricketts, C. (1996). Moult in relation to some aspects of reproduction and growth in swarms of Antarctic krill, Euphausia superba. Marine Biology, 127(2), 201–208. doi: 10.1007/BF00942104

48. Tarling, G. A., & Cuzin-Roudy, J. (2003). Synchronization in the molting and spawning activity of northern krill (Meganyctiphanes norvegica) and its effect on recruitment. Limnology and Oceanography, 48(5), 2020–2033. doi: 10.4319/lo.2003.48.5.2020

49. Hering, L, Henze, M.J., Kohler, M., Kelber, A., Bleidorn, C., Leschke, M., Nickel, B., Meyer, M., Kircher, M., Sunnucks, P., Mayer, G. (2012) Opsins in onychophora (velvet worms) suggest a single origin and subsequent diversification of visual pigments in arthropods. Molecular Biology & Evolution, 29(11), 3451–3458. doi: 10.1093/molbev/mss148

50. Hering, L., and Mayer, G. (2014). Analysis of the opsin repertoire in the tardigrade Hypsibius dujardini provides insights into the evolution of opsin genes in Panarthropoda. Genome Biol. Evol. 6, 2380–2391. doi: 10.1093/gbe/evu193

51. Colbourne, J. K., Pfrender, M. E., Gilbert, D., Thomas, W. K., Tucker, A., Oakley, T. H., et al. (2011). The ecoresponsive genome of *Daphnia pulex*. Science 331, 555–561. doi: 10.1126/science.1197761

52. Eriksson, B. J., Fredman, D., Steiner, G., and Schmid, A. (2013). Characterisation and localisation of the opsin protein repertoire in the brain and retinas of a spider and an onychophoran. BMC Evol. Biol. 13:186. doi: 10.1186/1471-2148-13-186

53. Futahashi, R., Kawahara-Miki, R., Kinoshita, M., Yoshitake, K., Yajima, S., Arikawa, K., et al. (2015). Extraordinary diversity of visual opsin genes in dragonflies. Proc. Natl. Acad. Sci. U.S.A. 112, E1247–E1256. doi: 10.1073/pnas.1424670112

54. Beckmann, H., Hering, L., Henze, M. J., Kelber, A., Stevenson, P. A., & Mayer, G. (2015). Spectral sensitivity in Onychophora (velvet worms) revealed by electroretinograms, phototactic behaviour and opsin gene expression. Journal of Experimental Biology, 218(6), 915–922. doi: 10.1242/jeb.116780

