## Supplementary material for "The most comprehensive annotation of the Krill transcriptome provides new insights for the study of physiological processes and environmental adaptation": Suppermentary Material

**Table S1. RNA-seq sample collection.** All samples are divided into groups according to the experimental settings: samples from Group 1 represent larval specimens; samples from Group 2 represent adult specimens and were described by Höring et al (2021); samples from Group 3 correspond to adult specimens coming from the RNA sequencing experiment available at the NCBI BioProject PRJNA640244; samples from Group 4 represent adult individuals and were described by Suter et al. [36]. Each sample is reported with its SRA and BioProject code. The other columns in the table indicate the library type used and the different experimental conditions.

| SRA | BioProject | Library | Condition | Season | Geographical Area | Developmental Stage | Sex | Tissue | Group |
|---|---|---|---|---|---|---|---|---|---|
| SRR5187773 | PRJNA362526 | Unstranded | Control $CO_2$ concentration | | | Larval - Calyptopis | | | Group 1 |
| SRR5187772 | PRJNA362526 | Unstranded | 1000 ppm CO2 concentration | | | Larval - Calyptopis | | | Group 1 |
| SRR5187771 | PRJNA362526 | Unstranded | 2000 ppm CO2 concentration | | | Larval - Calyptopis | | | Group 1 |
| SRR5187770 | PRJNA362526 | Unstranded | Control CO2 concentration | | | Larval - Furcilia | | | Group 1 |
| SRR5187769 | PRJNA362526 | Unstranded | 1000 ppm CO2 concentration | | | Larval - Furcilia | | | Group 1 |
| SRR5187768 | PRJNA362526 | Unstranded | 2000 ppm CO2 concentration | | | Larval - Furcilia | | | Group 1 |
| ERS2955456 | PRJEB30084 | Stranded | | Summer | Lazarev Sea | Adult | Female | | Group 2 |
| ERS2955458 | PRJEB30084 | Stranded | | Summer | Lazarev Sea | Adult | Male | | Group 2 |
| ERS2955460 | PRJEB30084 | Stranded | | Summer | Lazarev Sea | Adult | Female | | Group 2 |
| ERS2955461 | PRJEB30084 | Stranded | | Summer | Lazarev Sea | Adult | Female | | Group 2 |
| ERS2955462 | PRJEB30084 | Stranded | | Summer | Lazarev Sea | Adult | Female | | Group 2 |
| ERS2955457 | PRJEB30084 | Stranded | | Summer | Lazarev Sea | Adult | Male | | Group 2 |
| ERS2955459 | PRJEB30084 | Stranded | | Summer | Lazarev Sea | Adult | Male | | Group 2 |
| ERS2955465 | PRJEB30084 | Stranded | | Winter | Lazarev Sea | Adult | Female | | Group 2 |
| ERS2955466 | PRJEB30084 | Stranded | | Winter | Lazarev Sea | Adult | Male | | Group 2 |
| ERS2955467 | PRJEB30084 | Stranded | | Winter | Lazarev Sea | Adult | Male | | Group 2 |
| ERS2955468 | PRJEB30084 | Stranded | | Winter | Lazarev Sea | Adult | Male | | Group 2 |
| ERS2955463 | PRJEB30084 | Stranded | | Winter | Lazarev Sea | Adult | Female | | Group 2 |
| ERS2955464 | PRJEB30084 | Stranded | | Winter | Lazarev Sea | Adult | Female | | Group 2 |
| ERS2955480 | PRJEB30084 | Stranded | | Summer | South Orkney | Adult | Female | | Group 2 |
| ERS2955481 | PRJEB30084 | Stranded | | Summer | South Orkney | Adult | Female | | Group 2 |
| ERS2955482 | PRJEB30084 | Stranded | | Summer | South Orkney | Adult | Male | | Group 2 |
| ERS2955483 | PRJEB30084 | Stranded | | Summer | South Orkney | Adult | Male | | Group 2 |
| ERS2955484 | PRJEB30084 | Stranded | | Summer | South Orkney | Adult | Male | | Group 2 |
| ERS2955485 | PRJEB30084 | Stranded | | Summer | South Orkney | Adult | Female | | Group 2 |
| ERS2955450 | PRJEB30084 | Stranded | | Winter | Bransfield Strait | Adult | Female | | Group 2 |
| ERS2955451 | PRJEB30084 | Stranded | | Winter | Bransfield Strait | Adult | Male | | Group 2 |
| ERS2955452 | PRJEB30084 | Stranded | | Winter | Bransfield Strait | Adult | Male | | Group 2 |
| ERS2955453 | PRJEB30084 | Stranded | | Winter | Bransfield Strait | Adult | Male | | Group 2 |
| ERS2955454 | PRJEB30084 | Stranded | | Winter | Bransfield Strait | Adult | Female | | Group 2 |
| ERS2955455 | PRJEB30084 | Stranded | | Winter | Bransfield Strait | Adult | Female | | Group 2 |
| ERS2955469 | PRJEB30084 | Stranded | | Summer | South Georgia | Adult | Female | | Group 2 |
| ERS2955470 | PRJEB30084 | Stranded | | Summer | South Georgia | Adult | Male | | Group 2 |
| ERS2955471 | PRJEB30084 | Stranded | | Summer | South Georgia | Adult | Male | | Group 2 |
| ERS2955472 | PRJEB30084 | Stranded | | Summer | South Georgia | Adult | Female | | Group 2 |
| ERS2955473 | PRJEB30084 | Stranded | | Summer | South Georgia | Adult | Male | | Group 2 |
| ERS2955474 | PRJEB30084 | Stranded | | Summer | South Georgia | Adult | Female | | Group 2 |
| ERS2955475 | PRJEB30084 | Stranded | | Winter | South Georgia | Adult | Male | | Group 2 |
| ERS2955476 | PRJEB30084 | Stranded | | Winter | South Georgia | Adult | Male | | Group 2 |
| ERS2955477 | PRJEB30084 | Stranded | | Winter | South Georgia | Adult | Male | | Group 2 |
| ERS2955478 | PRJEB30084 | Stranded | | Winter | South Georgia | Adult | Male | | Group 2 |
| ERS2955479 | PRJEB30084 | Stranded | | Winter | South Georgia | Adult | Male | | Group 2 |
| SRR12042644 | PRJNA640244 | | Low temperature | | | | | | Group 3 |
| SRR12042643 | PRJNA640244 | | Low temperature | | | | | | Group 3 |
| SRR12042642 | PRJNA640244 | | Low temperature | | | | | | Group 3 |
| SRR12042641 | PRJNA640244 | | Mid-temperature | | | | | | Group 3 |
| SRR12042640 | PRJNA640244 | | Mid-temperature | | | | | | Group 3 |
| SRR12042639 | PRJNA640244 | | Mid-temperature | | | | | | Group 3 |
| SRR12042638 | PRJNA640244 | | High temperature | | | | | | Group 3 |
| SRR12042637 | PRJNA640244 | | High temperature | | | | | | Group 3 |
| SRR12042636 | PRJNA640244 | | High temperature | | | | | | Group 3 |
| Krill_Male_sex_a | AAS_4015_AAS_4015_Krill_Gonad_Transriptome* | Unstranded | | | | Adult | Male | Testis and tails | Group 4 |
| Krill_Male_sex_b | AAS_4015_AAS_4015_Krill_Gonad_Transriptome* | Unstranded | | | | Adult | Male | Testis and tails | Group 4 |
| Krill_Female_sex_a | AAS_4015_AAS_4015_Krill_Gonad_Transriptome* | Unstranded | | | | Adult | Female | Ovaries and tails | Group 4 |
| Krill_Female_sex_b | AAS_4015_AAS_4015_Krill_Gonad_Transriptome* | Unstranded | | | | Adult | Female | Ovaries and tails | Group 4 |
| Krill_Male_tissue_a | AAS_4015_AAS_4015_Krill_Gonad_Transriptome* | Unstranded | | | | Adult | Male | Testis | Group 4 |
| Krill_Male_tissue_b | AAS_4015_AAS_4015_Krill_Gonad_Transriptome* | Unstranded | | | | Adult | Male | Testis | Group 4 |

*Downloadable at https://data.aad.gov.au/metadata/records/fulldisplay/AAS_4015_Krill_Gonad_Transcriptome

**Fig S1. Phylogenetic relationships of *Euphausia superba* opsins shown as rectangular phylogram.** Scale bars indicate amino acid substitutions per site.

**Table S2. Command line for each *de novo* assembly reconstruction performed.** The column "Sample Group" refers to sample annotation as reported in Table S1. The command line for IDBA-tran program was the same for both sample groups as this software does not provide any option regarding the library type.

| Assembler | Library | Sample Group | Command Line |
|---|---|---|---|
| Trinity | Stranded | Group 2 | Trinity --seqType *fq* --max_memory *480G* --left <*reads_1.fastq*> --right <*reads_2.fastq*> --output <*output_dir*> --SS_lib_type *RF* --CPU *32* --monitoring |
| Trinity | Unstrandec | Group 1 | Trinity --seqType *fq* --max_memory *480G* --left <*reads_1.fastq*> --right <*reads_2.fastq*> --output <*output_dir*> --CPU *32* --monitoring |
| BinPacker | Stranded | Group 2 | BinPacker -d -s *fq* -p *pair* -l <*reads_1.fastq*> -r <*reads_2.fastq*> -g *200* -m *RF* -k *25* -o <*output_dir*> |
| BinPacker | Unstrandec | Group 1 | BinPacker -d -s *fq* -p *pair* -l <*reads_1.fastq*> -r <*reads_2.fastq*> -g *200* -k *25* -o <*output_dir*> |
| IDBA-tran | Stranded | Group 2 | idba_tran -r <*read.fasta*> -o <*output_dir*> --min_transcript *200* |
| IDBA-tran | Unstrandec | Group 1 | idba_tran -r <*read.fasta*> -o <*output_dir*> --min_transcript *200* |
| rnaSPAdes | Stranded | Group 2 | rnaspades.py -1 <*reads_1.fastq*> -2 <*reads_2.fastq*> --threads *32* --memory *490* -o <*output_dir*> --ss-rf |
| rnaSPAdes | Unstrandec | Group 1 | rnaspades.py -1 <*reads_1.fastq*> -2 <*reads_2.fastq*> --threads *32* --memory *490* -o <*output_dir*> |
| TransABySS | Stranded | Group 2 | transabyss --pe <*reads_1.fastq*> <*reads_2.fastq*> --SS --outdir <*output_dir*> --name <*output_name*> --length *200* |
| TransABySS | Unstrandec | Group 1 | transabyss --pe <*reads_1.fastq*> <*reads_2.fastq*> --outdir <*output_dir*> --name <*output_name*> --length *200* |

**Table S3. List of the 14 *Euphausia superba* putative opsins.**

| | Krill gene ID | Krill transcript ID | Gene name | % Identity | E-Value | Annotation | |
|---|---|---|---|---|---|---|---|
| Previously identified | ESG045640 | ESS176721 | EsRh1a | 96 | 0 | Euphausia superba opsin 1a | De Pittà *et al*. 2013 |
| | ESG047606 | ESS182754 | EsRh1b | 96 | 0 | Euphausia superba opsin 1b | De Pittà *et al*. 2013 |
| | ESG047724 | ESS183135 | EsRh2 | 98 | 0 | Euphausia superba opsin 2 | gi\|1004170845 |
| | ESG046593 | ESS179523 | EsRh3 | 93 | 0 | Euphausia superba opsin 3 | gi\|1004170847 |
| | ESG048023 | ESS1839951 | EsRh4 | 98 | 0 | Euphausia superba opsin 4 | gi\|1004170849 |
| | ESG047639 | ESS182855 | EsRh5 | 91 | 0 | Euphausia superba opsin 5 | gi\|1004170851 |
| | ESG047628 | ESS182813 | EsRh6 | 99 | 0 | Euphausia superba opsin 6 | gi\|1004170853 |
| | ESG050629 | ESS190453 | EsPeropsin | 98 | 0 | Euphausia superba peropsin | gi\|1004170855 |
| Putative new opsin | ESG045818 | ESS177277 | EsRh7 | 71 | 0 | Euphausia superba opsin 1b | De Pittà *et al*. 2013 |
| | ESG047527 | ESS182477 | EsRh8 | 59 | 1E-158 | Euphausia superba opsin 4 | gi\|1004170849 |
| | ESG047718 | ESS183103 | EsRh9 | 87 | 0 | Euphausia superba opsin 5 | gi\|1004170851 |
| | ESG047838 | ESS183478 | EsRh10 | 74 | 0 | Euphausia superba opsin 5 | gi\|1004170851 |
| | ESG044027 | ESS171537 | EsOnychopsin | 70 | 5E-151 | Penaeus vannamei onychopsin | ROT82196.1 |
| | ESG060540 | ESS214119 | EsArthropsin | 59 | 2E-49 | Penaeus vannamei putative rhodopsin-like | ROT82707.1 |

**Table S4. List of krill putative microRNAs precursors with corresponding known mature sequence and annotated GO terms.**
