## Supplementary figures and images for "The most comprehensive annotation of the Krill transcriptome provides new insights for the study of physiological processes and environmental adaptation"

### Figure S1

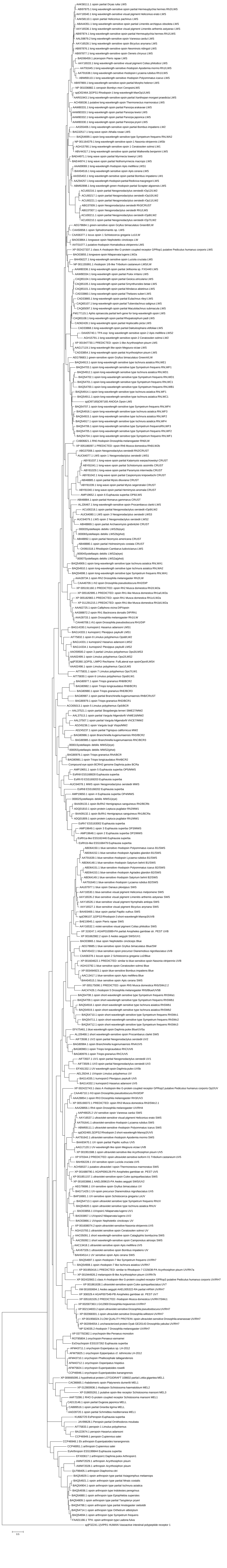
